# Artificial neural network filters for enhancing 3D optical microscopy images of neurites

**DOI:** 10.1101/441071

**Authors:** Shih-Luen Wang, Seyed M. M. Kahaki, Armen Stepanyants

## Abstract

The ability to extract accurate morphology of labeled neurons from microscopy images is crucial for mapping brain connectivity and for understanding changes in connectivity that underlie learning and memory formation. There are, however, two problems, specific to optical microscopy imaging of neurons, which make accurate neuron tracing exceedingly challenging: (*i*) neurites can appear broken due to inhomogeneous labeling and (*ii*) neurites can appear fused in 3D due to limited resolution. Here, we propose and evaluate several artificial neural network (NN) architectures and conventional image enhancement filters with the aim of solving both problems. To evaluate the effects of filtering, we examine the following four image quality metrics: normalized intensity in the cross-over regions between neurites, radius of neurites, coefficient of variation of intensity along neurites, and local background to neurite intensity ratio. Our results show that NN filters, trained on optimized semi-manual traces of neurites, can significantly outperform conventional filters. In particular, U-Net based filtering can virtually eliminate background intensity, while also reducing radius of neurites by 23% to nearly 1 voxel, decreasing intensity in the cross-over regions between neurites by 22%, and reducing variations in intensity along neurites by 26%. These results suggest that including a NN filtering step, which does not require much extra time or computing power, can be beneficial for neuron tracing projects.

## INTRODUCTION

Recent advances in genetic engineering and optical microscopy have allowed neuroscientists to label sparse populations of neurons and image their arbors in 3D on the scale of the entire mouse brain. At present, semi-manual tracing is the only reliable way of extracting information about the layout of axonal and dendritic arbors of individual neurons from such imaging data. However, semi-manual tracing methods are very time consuming and are prone to errors and user biases. Therefore, they are unsuitable for high-throughput neuron tracing projects. Inadequate image quality is the main obstacle on the way to accurate automated neuron tracing. NN image enhancement filters developed in this study can make imaging data more amenable to automated tracing.

## METHODS

We evaluated the effects of 3 NN filters (Figure 1) and 3 conventional filters on the quality of neuron images. The first NN filter is a shallow dense network with 1 hidden layer of 100 sigmoid units. It receives an input in the form of a 21×21×7 voxel sub-image and produces a single output that represents the enhanced intensity of the central voxel. The second network is a multilayer dense network with 3 hidden layers containing 100, 50, and 100 rectified linear units (ReLu) and sigmoid units in the output layer. It receives a 28×28×10 voxel sub-image as an input and produces an output in the form of an 8×8×4 voxel sub-image. The third network is a U-Net [1] with a 32×32×8 voxel input and output. It includes two dropout layers with dropout rates of 20%. Conventional 3D filters used in this study include the Laplacian of Gaussian (LoG) filter of NCTracer [2,3] (size is 2×2×2 voxels), the median filter of NCTracer (size is 3×3×3 voxels), and the MeanShift filter of ImageJ (spatial radius is 3 and color distance is 25). Parameters of conventional filters were tuned to the best of our ability to maximize image quality, and the resulting filter sizes roughly match the average width of neurites in the images.

**Figure 1:**
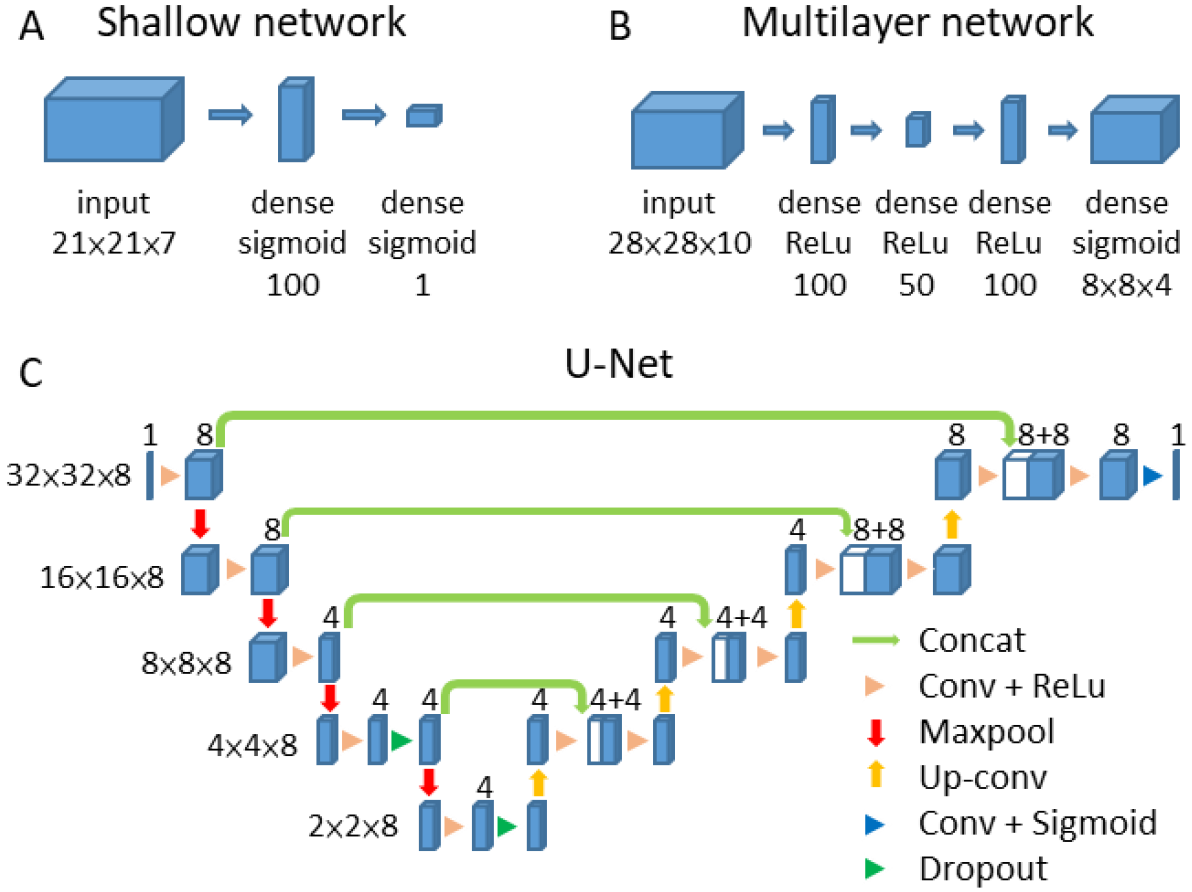
Architectures of the 3D NN filters used in this study. **A.** Shallow feedforward network with dense connectivity and sigmoidal neurons. Blue boxes represent neuron layers with neuron numbers shown. The network receives a 21×21×7 voxel sub-image as an input and generates a scalar output. **B.** Multilayer feedforward network with dense connectivity and ReLu neurons. This network has a bottleneck. It receives a 28×28×10 voxel sub-image as an input and generates an 8×8×4 voxel output. **C.** U-Net [1]. Blue boxes represent feature maps with the number of channels denoted above each box. The network receives a 32×32×8 voxel input and generates an output of the same size.

All filters were tested on the dataset of Neocortical Layer 1 Axons [4] used in the DIADEM challenge [5]. This dataset includes 6 stacks of two-photon microscopy images containing fluorescently labeled axons (Figure 2). The image stacks consist of 33-60 planes, each of which is 512×512 voxels in size. All axons were semi-manually traced and optimized in the NCTracer software. Optimized traces were used to create labeled images in which voxels at distances of <1 voxel size away from the trace are labeled as foreground, 1 voxel size away as undetermined (not used for training), and >1 voxel size away as background. Four image stacks were used for training NN filters, one was used for validation, and one for testing.

## RESULTS

Figure 2 shows the maximum intensity projection of one of the image stacks along with the outputs of the 6 filters. We note that conventional metrics, such as precision and recall, are not well suited for the task of assessing the quality of neuron images. This is because such measures treat image voxels independently, without regard for their connectivity. Therefore, we developed 4 new metrics, which, in our opinion, capture morphological features that are essential for accurate neuron tracing. The first metric reflects the normalized intensity in cross-over regions formed by adjacent (within 10 voxels) axons. Such axons are often interconnected by tracing algorithms, and, therefore, having low intensity in cross-over regions is favorable for tracing. Specifically, this metric is defined as the ratio of average intensity along the shortest A* [6] path connecting the axons to the average intensity of the two axons in the vicinity of the cross-over (see Figure 3A inset and legend for details). The second metric is designed to reflect inhomogeneity of intensity along axons in a filtered image. It is defined as the coefficient of variation (CV) of intensity along the centerline of a neurite. Because abrupt changes in intensity can lead to broken neuron traces, low CVs are advantageous for tracing. The remaining two metrics are the mean neurite radius (measured in the NCTracer software) and local background to foreground intensity ratio as defined by the labels.

**Figure 2:**
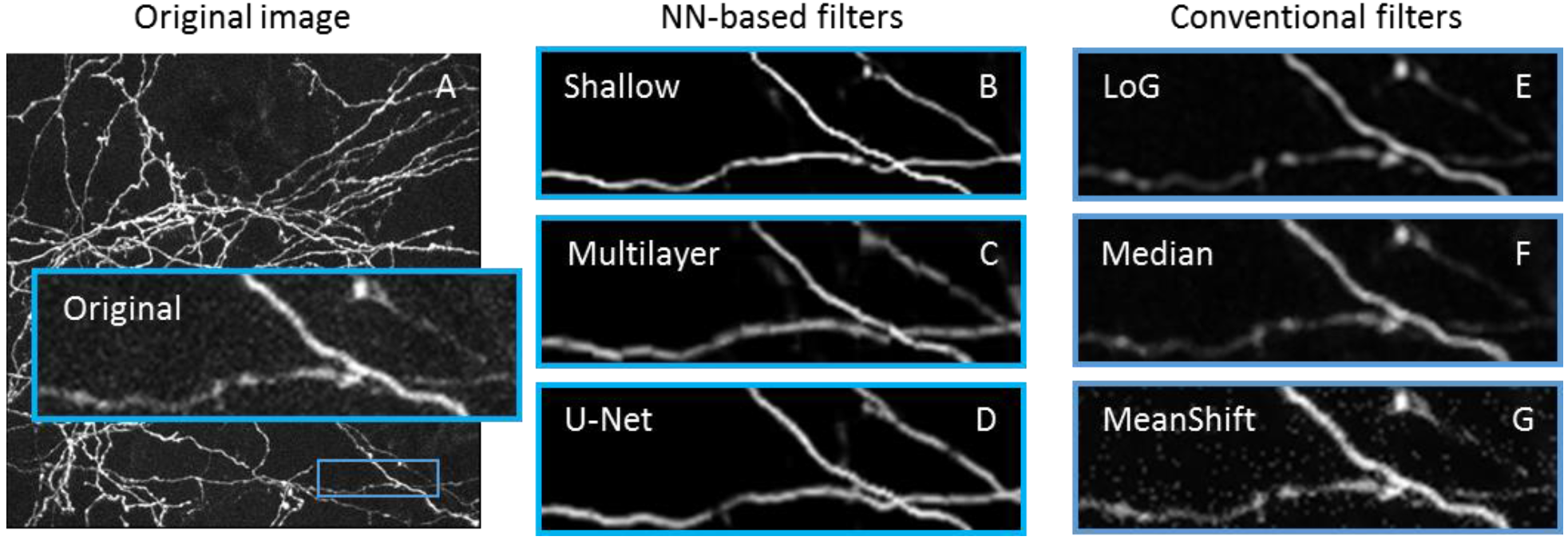
Images of neurites enhanced with the proposed and conventional image processing filters. **A.** Maximum intensity projection of an image stack containing layer 1 axons of mouse neocortical neurons [4]. Inset shows a 4× zoomed-in view of a small region outlined with the cyan rectangle. **B-G.** The same region in the images enhanced with NN (B-D) and conventional filters (E-G). The widths of the cyan rectangles are 40 μm.

Figure 3 shows the four metrics (mean ± s.e.m.) calculated based on the original and filtered images. NN filters can significantly improve the image quality, and generally outperform conventional filters. The U-Net filter produces particularly promising results (Figure 2B). It virtually eliminates background intensity, while also reducing neurite radius by 23% to nearly 1 voxel, decreasing intensity in the cross-over regions between neurites by 22%, and reducing variations in intensity along the neurites by 26% (Figure 3).

**Figure 3:**
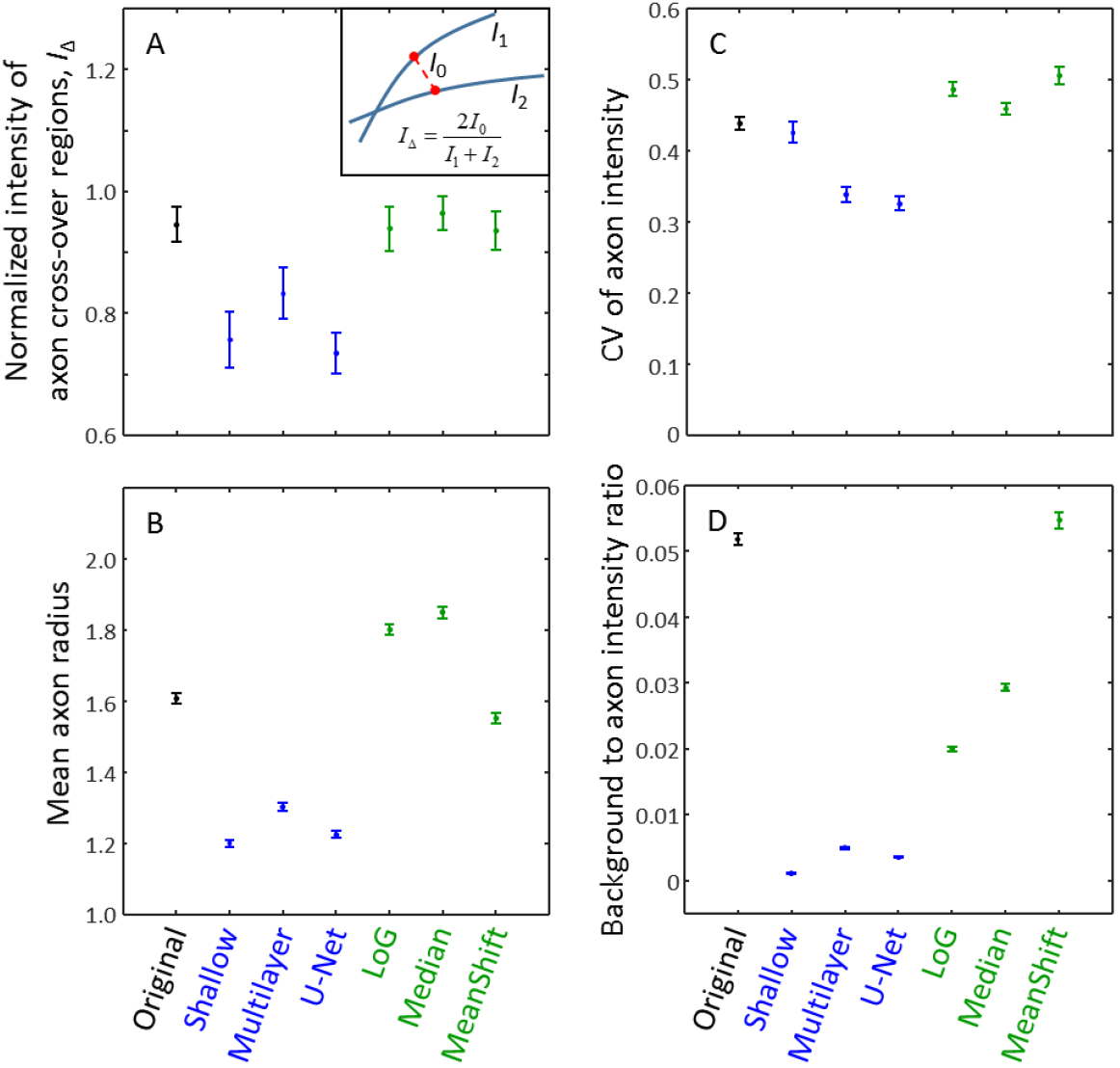
Quality of images enhanced with the NN (blue error bars) and conventional (green error bars) filters. **A.** Normalized intensity in cross-over regions between axons. Inset illustrates how the metric is calculated. Cross-over is defined as a location in the image stack where two axons come within 10 voxels of each other. *I*_1_ and *I*_2_ denote the average intensities of A* paths [6] along the axons in the vicinity (within 5 voxels) of the cross-over region. *I*_0_ is the average intensity of the shortest A* path connecting the two axons (red dashed line). **B.** Axon radius calculated for voxels on A* axon centerlines. **C.** Coefficient of variation of intensity on A* axon centerlines. **D.** Local background to foreground ratio. Local intensities are based on randomly sampled 64×64×10 sub-images, in which background and foreground voxels are defined by the label. Error bars correspond to s.e.m.

Even the shallow NN significantly outperforms conventional image processing filters. In comparison with U-Net, it produces a somewhat lower background intensity (Figure 3D) and slightly thinner axons (Figure 3B), which, however, comes at the expense of axon intensity variations (Figure 3C).

## CONCLUSION

We showed that NN filters can be successfully used to enhance morphological features of neurites in 3D optical microscopy images. Even the shallow NN outperforms conventional image processing filters in terms of the four morphology-related metrics introduced in this study. These metrics can be used to assess the appropriateness of different neuron labeling and imaging methods for neural tracing applications, including circuit mapping and structural plasticity studies. It remains to be seen if enhancements observed in the filtered images will be reflected in the accuracy of neuron traces.

NN image enhancement filters described in this study are freely available at https://github.com/neurogeometry/NNfilters. We thank Rohan Gala for helpful discussions. This work was supported by the NIH grant R01 NS091421.

